# Microscopic and Macroscopic Response of a Cortical Neuron to an External Electric Field Computed with the Boundary Element Fast Multipole Method

**DOI:** 10.1101/391060

**Authors:** Sergey N Makarov, Gregory M Noetscher, Padmavathi Sundaram

## Abstract

The goal of this study is to demonstrate how one can compute the activating function and surface charge density resulting from application of an external electric field to a high-resolution realistic neuronal morphology. We use the boundary element fast multipole method (BEM-FMM) on an ordinary computer to accurately perform these computations in under 2-10 minutes for a dense surface mesh of a single neuron with approximately 1.4 million triangles. Prior work used commercial finite element method (FEM) software which required creation of a volumetric tetrahedral mesh between fine neuronal arbor, potentially resulting in prohibitively large volume sizes and long mesh generation times. We used the example of a human pyramidal neuron with an externally applied *E*-field to show how our approach can quickly and accurately compute the induced surface charge density on the cell surface and the activating function of the cable equation. We found that the induced surface charge density perturbs the macroscopically applied *E*-field on a microscopic spatial scale. The strength of the perturbation depends on the conductivity contrast; the stronger the contrast, the larger the perturbation. In our example, the induced surface charge density may change the average activating function by up to 75%. We also embedded this neuron model into a detailed macroscopic human head model and simulated a realistic TMS excitation using the BEM-FMM method for the combined model. The solution obtained in this case predicted a smaller activating function error. The difference between the microscopic and the macroscopic effect of the externally applied electric field is of much interest to users of extracellular stimulation techniques and merits further study.

## 1. Introduction

Neuronal activity in the cortex can be modulated using many possible techniques such as transcranial magnetic stimulation (TMS), transcranial electric stimulation (TES), and intracortical microstimulation (ICMS). The effectiveness of any given method depends on which neuron populations are modulated by the stimulation given specific spatial and temporal parameters [1],[2]. Computational models can be used to better understand how a given neuron experiences the externally applied electric field as a function of its morphology, location and orientation.

There are many recent computational studies that simulate electrical stimulation of multicompartment neuron models with realistic morphologies. Seo et al. [3] modeled cortical pyramidal neurons with the activating function derived from an external TMS field, which was obtained based on an individual macroscopic head model. Goodwin & Butson [4] also modeled cortical neurons subject to an external TMS field. The subject-specific external TMS field was computed using realistic head and coil models and then applied to each neuron model to simulate its response based on the cable equation. Similarly, Aberra et al. [5] modeled cortical neurons with a detailed morphology subjected to an externally applied E-field which was projected onto the curved axonal and dendritic compartments of the neuron.

These recent efforts use the “hybrid FEM cable-equation approach” (see Joucla et al. [6] for a review). First, they accurately compute the external macroscopic electric field/potential at the neuron position using the finite element method (FEM) (or an analytical equation). Then, this computed field is directly applied to a neuronal morphology, described by a number of one-dimensional straight segments in space. This gives us a local activating function for the one-dimensional cable equation along the length of the neuron solved in NEURON software [7]. The activating function is proportional to the second spatial derivative of the extracellular electric field along the length of the neuron [8].

It is however known that the externally applied electric field will cause induced charges to reside on any physical interface (neuron membrane) separating two media with different conductivities. These charges will generate their own secondary field and electric potential, which will perturb the macroscopically applied electric field and may even affect the estimates of the above approach on a microscopic scale. It is therefore important to understand when the assumption of an unperturbed externally applied macroscopic stimulation field close to the neuron is applicable. In [9], the authors suggest that “this additional polarization typically has negligible amplitude and does not affect computational models.” Given the importance of this effect to users of extracellular stimulation technologies, this statement may need to be tested in realistic neuron models.

One challenge is that it is difficult to model detailed cortical neuron morphologies using FEM. Each neuronal compartment must be represented as a thin tubular surface comprised of a triangular or quadrilateral mesh with at least one shell. Creating a volumetric tetrahedral or hexahedral mesh between many curved thin branches of the neuronal arbor is a difficult technical task which may result in prohibitively large volumetric mesh generation times and extremely large volumetric mesh sizes. This is perhaps why many of the related prior FEM modeling efforts ([10],[11],[6]) are restricted to a smaller number of straight cylinders (either joined or not) within the COMSOL environment, a general-purpose commercial finite element software with a relatively low speed.

In this study, we show how one can compute an external electric field for a realistic neuron with dense morphology efficiently using an alternative dedicated numerical approach, the boundary element fast multipole method (BEM-FMM) [12]. This method is fully compatible with the MATLAB^®^ environment (Mathworks, Natick, MA). It is based on the surface-charge formulation of the BEM equations [13],[14],[15] and involves accurate computations of neighbor surface integrals [13] using an efficient FMMLIB3 library of a precompiled fast multipole method developed by Drs. Greengard and Gimbutas [16]. We have already applied the BEM-FMM method to TMS [12] and EEG/MEG related problems.

Here, we first considered the simple case of a single human pyramidal neuron in a homogeneous extracellular volume subjected to a constant external electric field. Our computations were based on the “hole model” (see [17],[18],[19]) which involves computing around the cell by modeling the intracellular space as a nonconducting region. Effectively, the intracellular conductivity equals zero, which allows computation of a steady state distribution, that accurately accounts for the cell presence effects on the field [17],[18],[19]. Along with the precise hole model, we also tested some intermediate finite values of the intracellular conductivity and the corresponding conductivity contrast. This was done to highlight the method convergence, better understand the results, and establish a bridge to the coupled dynamic neuron-field model which indeed uses finite interacellular conductivity values.

Our results suggest that the externally induced membrane charges may change the one-dimensional averaged activating function by up to 75% in contrast to the effect size suggested by the approaches used in the prior work cited above.

Finally, the detailed Computer Aided Design (CAD) neuron model with 1.4 M facets was embedded into the macroscopic CAD head model with 2.8 M facets (specifically, into a cortical sulcus). We assumed that the neuron was simply an extra brain compartment and, using the BEM-FMM method, ran coupled realistic multiscale TMS simulations for the head and the neuron as one single boundary element problem, without using any extra simplifications.

## 2. Materials and Methods

### 2.1. Neuron morphology

We used a digital morphological reconstruction of a human neocortical pyramidal neuron H16-03-001-01-09-01_559391771_m, ID NMO_86955 from the NeuroMorpho.Org inventory Version 7.5 [20]. The specific reconstruction contained a total of 81 segments: 23 axonal branches, 40 dendritic branches, 18 apical dendrite branches, and a (spherical, 6.3 µm radius) soma. The total number of individual straight geometrical segments or edges included in the morphology file was 9237 and the total number of nodal points was 9318. The neuron length was about 900 µm.

### 2.2. CAD neuron model

We developed an algorithm to create a high-quality strictly 2-manifold or watertight “tubular” triangular mesh of arbitrary resolution around an arbitrary-bent single fiber path. The implementation was in MATLAB^®^ (Mathworks, Natick, MA). The algorithm also creates a start and an end cap for the tubular mesh. We applied this algorithm to every segment of the neuron. Further, using Autodesk Meshmixer software (Autodesk, San Rafael, CA), the meshed segments were semi-manually interconnected to each other and to the soma so as to create single mesh surface. Finally, we applied a surface-preserving Laplacian smoothing process [21] to the final mesh. Fig. 1 shows the resulting high-quality triangular surface mesh. The mesh had 1.4 M triangles with the average mesh quality (defined as twice the ratio of incircle radius to the circumcircle radius of a triangle) of 0.64.

**Fig. 1.**
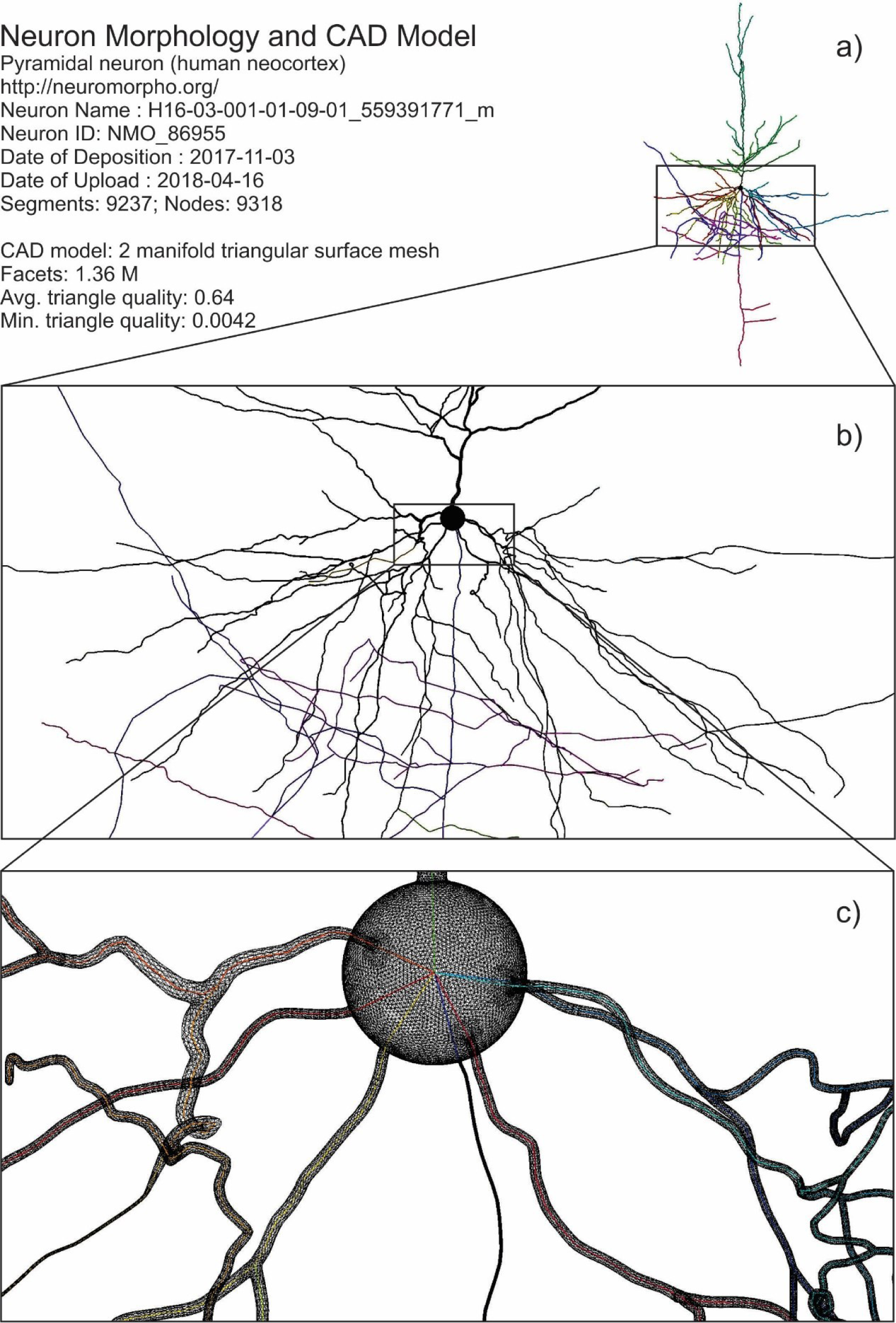
Neuron morphology a) and its CAD model b), c). The soma is modeled by a sphere with the given radius of 6.3 µm.

### 2.3. Boundary element fast multipole method

We used the fast multipole method or FMM introduced by Rokhlin [22] and Greengard and Rokhlin [23]. Conceptually, FMM is somewhat similar to the fast Fourier transform but developed for a complicated 3D morphology. It can speed up computation of a matrix-vector product by many orders of magnitude. The matrix-vector product naturally appears when the field from many point sources in space has to be computed at many observation points. This is exactly the task of the boundary element method or BEM. Makarov et al. recently developed an integration of the BEM and FMM for quasistatic electromagnetic computations [12] in an effort improve the BEM performance. The BEM-FMM method overcomes the major disadvantage of the BEM which is the inability to solve large high-resolution models, using a fast iterative algorithm and without system matrix storage. At the same time, the major advantage of the BEM – superior accuracy close to the boundaries and in the surounding space – is retained.

Specifically, we used the efficient precompiled FMMLIB3 library of the fast multipole method developed by Drs. Greengard and Gimbutas [16]. This library is fully compatible with the MATLAB environment (both Windows and Linux) and has already demonstrated excellent performance for TMS [12] and EEG/MEG related problems. We also used a native MATLAB generalized minimum residual (GMRES) iterative algorithm written by Drs. P. Quillen and Z. Hoffnung of MathWorks, Inc.

Fig. 2 shows the BEM-FMM algorithm workflow used in this work. The fast multipole method was applied three times: (i) for computing the primary fields (electric or magnetic or the corresponding potential(s)) at all conductivity boundaries and in space, (ii) for solving the BEM equation iteratively and in a matrix-free form, and (iii) for computing the secondary and the total electric and magnetic fields.

**Fig. 2.**
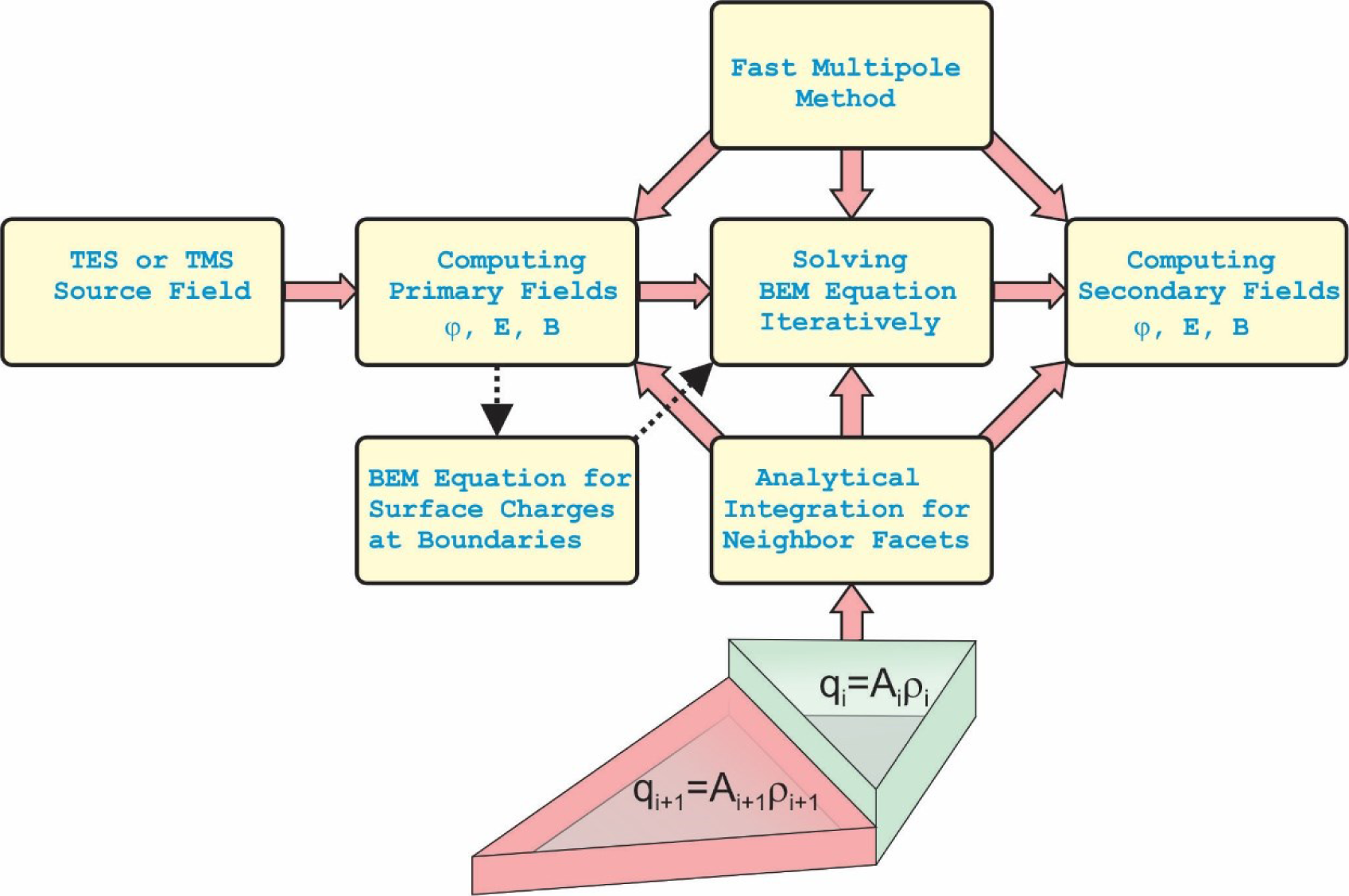
Flow chart of the fast multipole boundary element method (BEM-FMM) used in this study. Generally, the source field is due to the externally applied electric field injected by TMS or TES.

The major modification compared to our prior work [12] is that we accurately computed the primary and secondary electric potential with the fast multipole method. We also employed the generalized minimum residual method for the iterative solution itself. This method significantly reduces computational time. Finally, we explicitly enforced the charge conservation law, which is a must for accurate high-resolution computations of the electric potential.

Numerical routines that generated results of this study along with the neuron CAD model have been organized in the form of a self-consistent code which utilizes the widely accessible MATLAB platform for both Windows and Linux. This code will be made available through MATLAB Central.

### 2.4. Externally Applied E-field

In its simplest form, the externally applied electric source field, **E**^p^(**r**), is constant and is directed along the neuron pointing toward the axon,

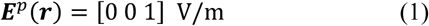

We assumed a value of 1 V/m for the *E*-field; since all results of extracellular computations given below are linear, they can be trivially scaled for different *E*-field strengths.

### 2.5. Material parameters

There are only two parameters for the present single-shell steady state neuronal model: the extracellular conductivity and the intracellular conductivity. Following Refs. [24],[17], we assigned the conductivity values around the numbers: *σ_e_* = 20 mS/m for the extracellular space and *σ_i_* = 5 mS/m for the intracellular space. We varied *σ_e_* and *σ_i_* according to Table 1 given below. This is done to investigate the effect of the conductivity contrast, 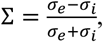, on the solution and the activating function, which will be an important result of this study.

**Table 1.** Convergence of the iterative BEM-FMM solution depending on the conductivity contrast. The maximum relative residual (the stopping criterion) is fixed at 1e-6 in every case. Time per one iteration is 9 sec.

| Setup # | $\sigma_e$ | $\sigma_i$ | Dimensionless conductivity contrast $\Sigma$ | # of iterations needed to converge | Solution run time, sec |
| --- | --- | --- | --- | --- | --- |
| 1 | 33 mS/m | 0 mS/m<br>(precise hole model) | 1 | 76 | 684 |
| 2 | 33 mS/m | 1 mS/m | 0.94 | 45 | 405 |
| 3 | 18 mS/m | 2 mS/m | 0.80 | 20 | 180 |
| 4 | 20 mS/m | 5 mS/m | 0.60 | 13 | 117 |

## 3. Results

### 3.1. Method convergence and speed

Table 1 reports method convergence (the maximum relative residual of the BEM system of equations is fixed at 1e-6) for four different values of the conductivity contrast, Σ = 0.6, 0.8, 0.94, and 1.00. The convergence rate was promising, but it also showed a slow-down for larger contrast values. The solution run times are given for an Intel Xeon E5-2698 v4 CPU (2.20 GHz) Windows Server and for MATLAB^®^ 8.2 platform. Similar results have been obtained for the maximum relative residual of 1e-9 and for the intermediate values of the residual. We also computed the relative least squares error of the iterative solution for the surface charge density itself. We found that, at the end of the iteration process, this error did not exceed 1e-6 – 1e-7. All computations in this study were performed using a single processor.

### 3.2. Induced charge density on the membrane surface and additional extracellular potential

Fig. 3 shows the induced charge density in C/m^2^ due to the external applied electric field (Eq. 1) on the surface of the neuron. This charge density resides on the surface separating two media with different conductivities. The results in Fig. 3a and Fig. 3b are given for the conductivity contrast Σ = 1.00 and Σ = 0.60, respectively. In both cases, we found that the surface charge was distributed such that the electric field due to this surface charge tries to cancel the primary or external electric field inside the neuron compartments. We found that this dipole-like local distribution across the fiber was significantly stronger for the large conductivity contrast value (Fig. 3a) and less profound for the weaker conductivity contrast (Fig. 3b).

**Fig. 3.**
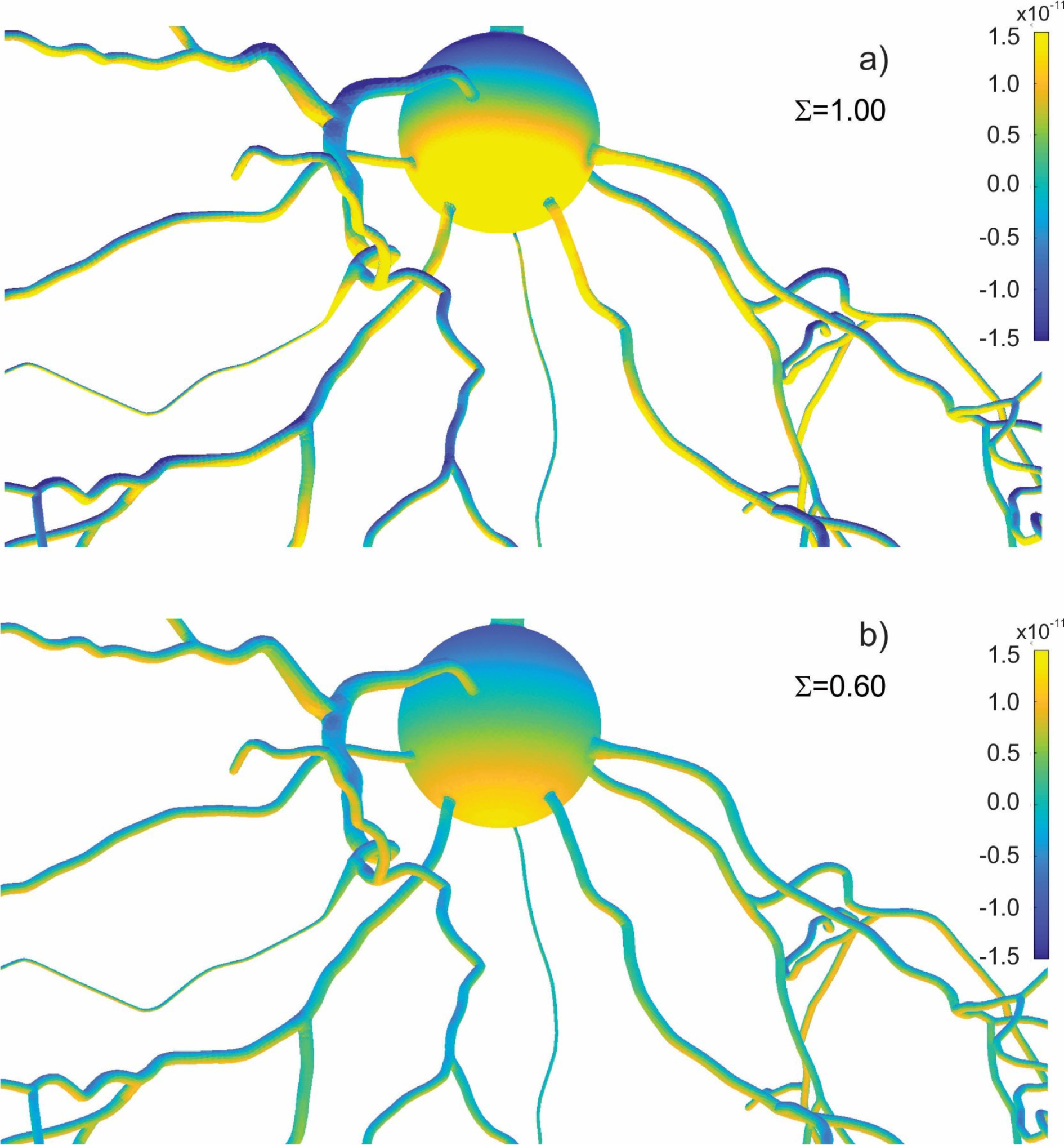
Induced surface charge density in C/m^2^ due to the external applied electric field (Eq. 1) on the surface of the neuron. The results in Fig. 3a are given for the conductivity contrast Σ = 1.00 and in Fig. 3b for the conductivity contrast Σ = 0.60.

The induced charge distribution gives rise to an extra electric potential that alters the extracellular potential values obtained directly from the primary field without the cell. This contribution was found to be rather small as compared to the primary potential for all three cases considered. Fig. 4 shows the induced or additional potential distribution in volts due to the external applied electric field (Eq. 1) exactly on the surface of the neuron. The results in Fig. 4a and Fig. 4b which again correspond to Σ = 1.00 and Σ = 0.60, respectively. We note that the induced potential no longer has a dominant dipole-like distribution across the fiber; there is also significant variation along the fiber.

**Fig. 4.**
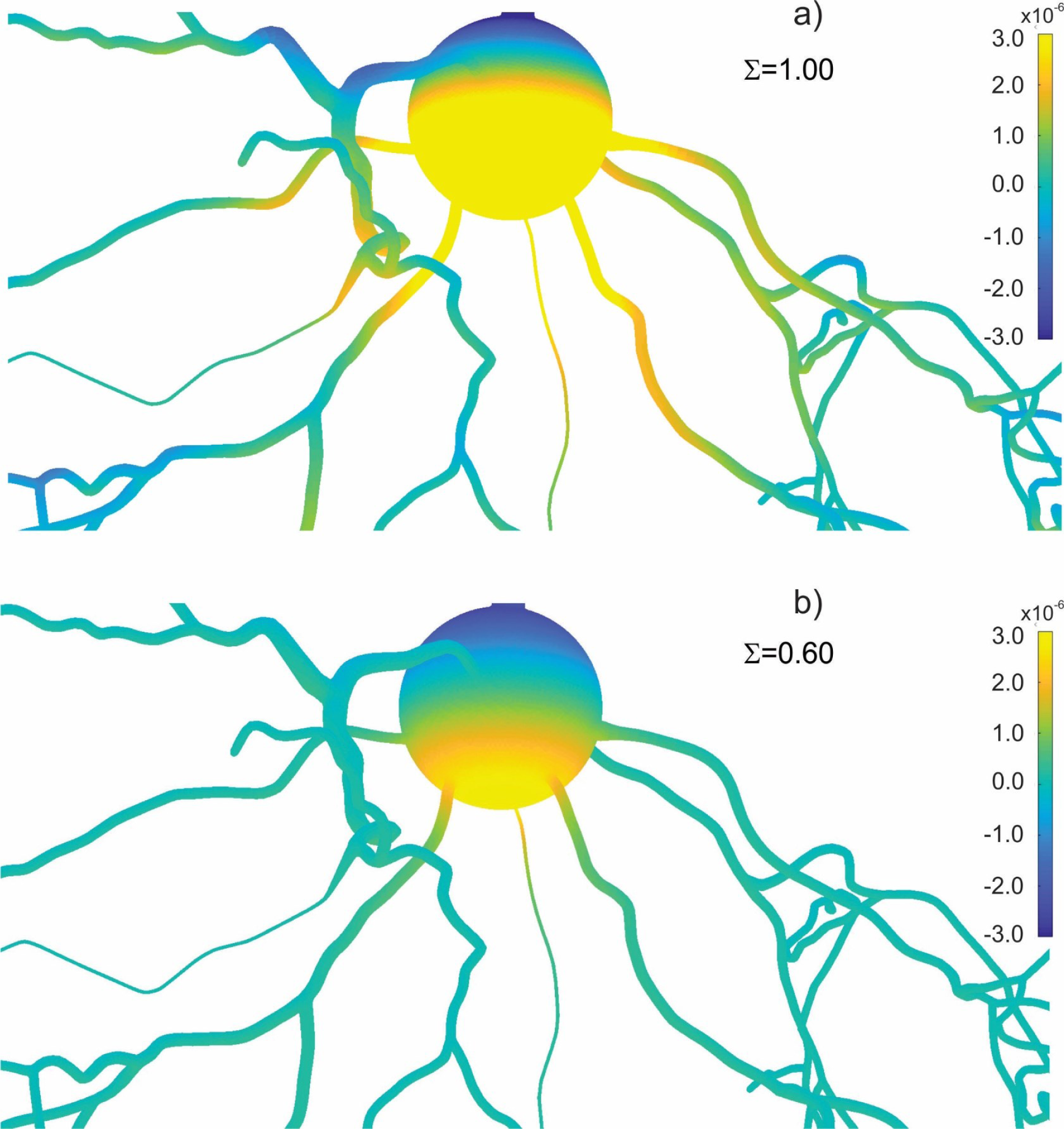
Induced (or secondary) electric potential in V due to the surface charge on the surface of the neuron. The results in Fig. 3a are given for the conductivity contrast Σ = 1.00 and in Fig. 3b for the conductivity contrast Σ = 0.60.

### 3.3. Effect of induced charges and induced potential on total extracellular potential

While the induced surface charge is not directly coupled to the cable equation, the extracellular potential is directly coupled. The one-dimensional neuron model based on the cable equation requires that the *average* value of the total extracellular potential 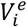 [25] be defined for every node *i* of the centerline of every fiber in the neuron model. In order to do so, we (i) find all triangular facets *j* which are closer to node *i* than to any other node, (ii) compute the potential value 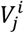 for every such facet and (iii) average the results with the weights inversely proportional to the respective distances,

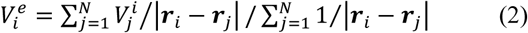

This method works well for a mesh of nearly equilateral triangles, i.e. for the present model. If necessary, weighting by triangle areas can be added.

Figs. 5 a, b demonstrate the distribution of the extracellular potential (Eq. 2) along dendrite #1. The *x*-axis is the distance along the dendrite. The blue curve is the primary extracellular potential, which was obtained without the charge density induced on the membrane surface and directly follows Eq. (1) after subsequent integration and projection onto the fiber path.

**Fig. 5.**
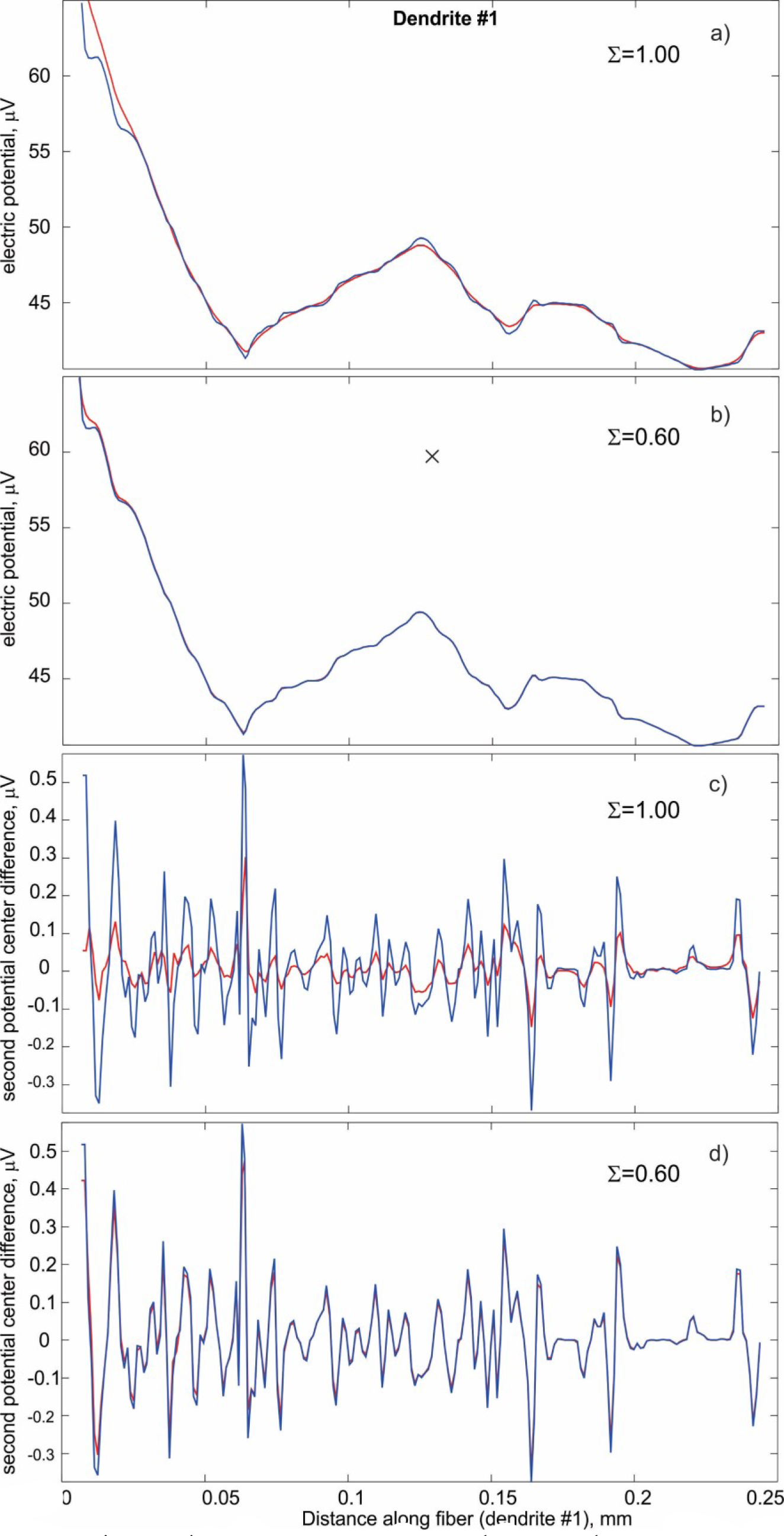
(a, b) – Total electric potential; (c, d) – its second center difference along centerline of dendrite #1. Blue curves: solution without surface charges; red curves – with surface charges.

The red curve is the total extracellular potential, which takes into account the contribution of induced surface charges. The results in Fig. 5a and Fig. 5b are given for the conductivity contrast Σ = 1.00 and Σ = 0.60, respectively.

In both figures, the difference between the two potential curves is quite small. However, we observe that the additional potential contribution attempts to significantly *smooth* the resulting potential curve and make it less jumpy. Those jumps are originally due to the complicated path shape of the dendrite itself. The smoothing effect is better seen in Fig. 5a; it is equivalent to reducing the second derivative of the corresponding potential function. We observed a similar smoothing effect for all other dendrites and axons. The effect was weaker for the lower conductivity contrast as Fig. 5b shows.

### 3.4. Effect of induced charges and induced potential on activating function

Although Figs. 5 a, b indicate only a small deviation for the potential itself, the result changes significantly when the second derivative of the extracellular potential is considered. To within a constant, this is exactly the activating function of the cable equation. The second derivative is proportional to the central difference,, [26] given by

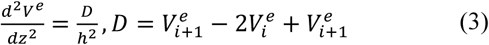

where is the distance along the path. Figs. 5 c, d plot the central difference *D* given by Eq. (3) in volts. We ignored the constant discretization step *h*, which does not add significance to the result. The blue curve is the second extracellular potential derivative without the induced charges. The red curve is the second extracellular potential derivative, which takes into account the contribution of induced surface charges. The results in Fig. 5c and Fig. 5d are given for the conductivity contrast Σ = 1.00 and Σ = 0.60, respectively. The key observation is that the exact electrical solution makes the activating function significantly more smooth due to the effect of the induced charges. Indeed, this only happens for the large conductivity contrast at the membrane as in Fig. 5c.

### 3.5. Activating function error

Finally in this section, we present in Fig. 6 the least squares error in the activating function for all dendrites and axons of the neuron. This error is the relative difference between the solution without the induced charges and the solution with the induced charges. When the conductivity contrast is high as in Fig. 6a, the average error may be on the order of 75%. When the conductivity contrast is relatively low as in Fig. 6d, the average error approaches 10% or so and becomes reasonably small as suggested in [9].

**Fig. 6.**
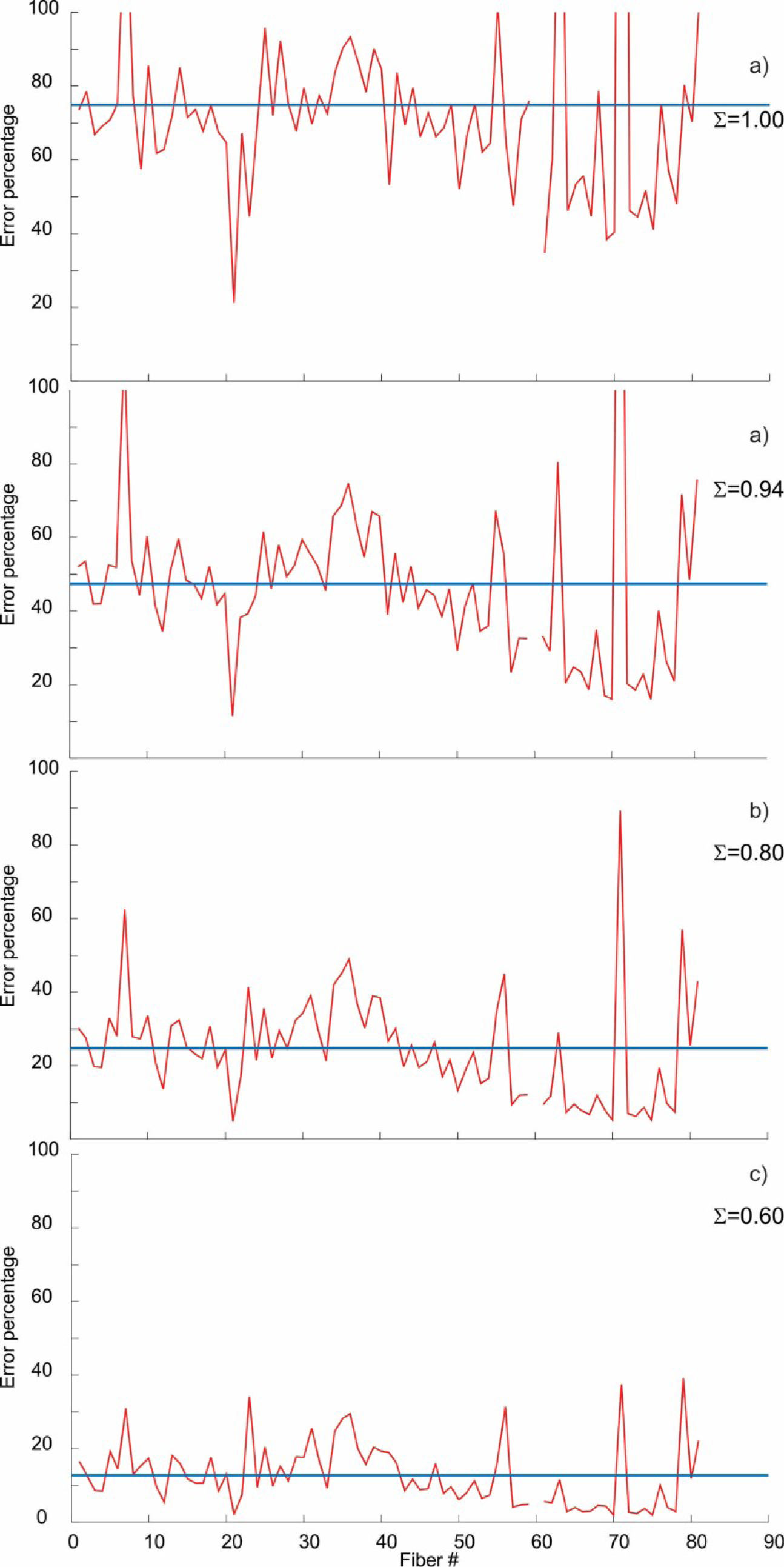
Least squares error in the activating function for all dendrites and axons of the neuron as compared to the no-induced charge solution. Horizontal line shows the mean value.

### 3.6 Application example

We attempted to compute the activating function for a realistic TMS scenario. A surface head model corresponding to subject #101309 from the Population Head Model Repository [27],[28] and the Connectome Project [29] was used as a starting point. The original model included the following seven compartments: cerebellum, CSF, GM, skin (or scalp), skull, ventricles, and WM, and results in 0.7 M triangular facets in total. The average edge length is 1.5 mm.

We refined the original surface meshes using a 1 × 4 barycentric triangle subdivision and then applied surface-preserving Laplacian smoothing [21]. The resulting mesh had an average resolution of 0.7 mm (0.6 mm for GM and WM) and 2.8 M triangular facets in total. Using the BEM-FMM solver, this macroscopic model alone ran reasonably fast (under 9 min in Linux MATLAB given 14 iterations and a relative residual below 10^−4^). Note that a finite element tetrahedral mesh for the same problem would result in at least 10 M volumetric elements as indicated by a test run with the commercial FEM software ANSYS Electronics Desktop Maxwell 18.2 2017. We assigned the following material conductivities: 0.126 S/m for cerebellum, 2.0 S/m for CSF, 0.106 S/m for GM, 0.333 S/m for skin (scalp), 0.0203 S/m for skull, 2.0 S/m for ventricles, and 0.065 S/m for WM. Grey matter and white matter shells of the head model are shown in Fig. 7a.

**Fig. 7.**
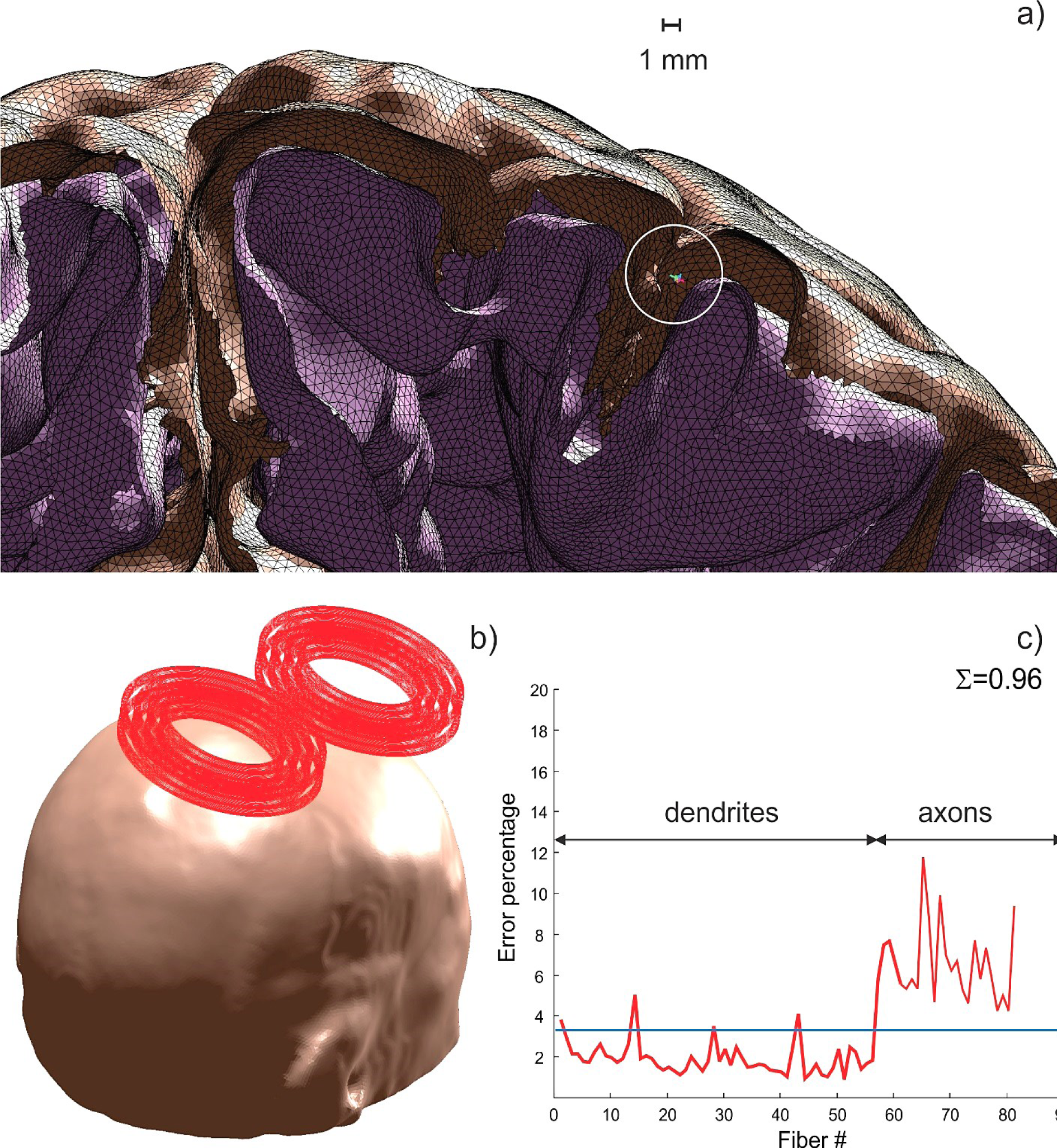
a) – Combined microscopic neuron and macroscopic tissue (grey and white matter) compartment model. Neuron position in a cortical sulcus is marked by a circle; (b) coil model for the coupled solution; c) least squares relative error in the activating function for all dendrites and axons of the neuron as compared to the no-induced charge solution.

The TMS coil model is the MRi-B91 of MagVenture, Denmark; this commercially available figure-of-eight coil with a rectangular conductor cross-section is shown in Fig. 7. To compute its incident field, we subdivided the coil conductors into about 100,000 small straight current elements and applied the fast multipole method again [12].

The neuron model from Fig. 1 with 1.4 M facets was embedded into the macroscopic CAD head model, specifically into a cortical sulcus as shown in Fig. 7a. The total combined model size was 4.2 M triangular faces. We assumed that the neuron was simply an extra brain compartment and ran the coupled multiscale simulations for the head and the neuron as one single boundary element problem, without using any additional asumptions. We assigned the intracellular conductivity as 2 mS/m so that the conductivity contrast with grey matter reaches Σ = 0.96. The coil position in Fig. 7b was chosen in such a way that the coil incident or primary unsteady electric field was aligned nearly parallel to the neuron. One iteration time for an Intel Xeon E5-2698 v4 CPU (2.20 GHz) Windows Server and for MATLAB 8.2 platform was 51 sec. However, many (about 100) such iterations were necessary to achieve a reasonable relative residual value of 1e-8. Again, the convergence varies depending on the value of the conductivity contrast.

Similar to Fig. 6, we present in Fig. 7c the least squares error in the activating function for all dendrites and axons of the neuron. This error is the relative difference between the solution without the induced charges and the solution with the induced charges. To obtain the former solution, the intracellular conductivity value was made equal to the grey matter conductivity value. From Fig. 7c, we observe that a small relative average error (about 2% for dendrites and 7% for axons) was generated by the surface charges; it appears to be larger for the axonal branches. This result is in a certain disagreement with the previous study concerned with the neuron alone (Fig. 6). However, this result has been verified for different values of the membrane conductivity and slightly different neuron positions. It may be possible that an “all in one” approach is not the best solution for the present coupled problem since the iterative solution accuracy may vary for the neuron and the large brain compartments, respectively, despite the overall small residual. To overcome this potential problem, we could extract the final field distribution from the macroscopic model and then solve the neuron alone. In this case, the presumably small effect of the neuron on the macroscopic cortical charge distribution can be ignored.

## 4. Discussion and Conclusions

In this study, the BEM-FMM method demonstrated an excellent convergence for the biophysically detailed single-neuron model subject to an externally applied electric field in terms of the extracellular problem statement. The relative residual values of 1e-9 and the relative iterative solution differences of 1e-7 have been achieved for all considered cases and for all conductivity contrasts in a relatively short amount of time. The convergence is fastest for smaller values of the conductivity contrast. Multiple and tightly spaced neurons could probably be modeled in a similar way. The BEM-FMM method does not have constraints on the model size or on the hard disk space although the RAM requirements may be demanding.

We employed the obtained solution in order to quantify the effect of the induced surface charge density on the activating function of the cable equation using the hole neuron model in the steady state. We found that, for sufficiently large conductivity contrasts (including the terminal value of 1) between the intracelluilar and the extracellular volume, the corresponding error in the activating function may be as large as 75%. Finally, we embedded the neuron model into a detailed macroscopic head model and simulated a realistic TMS excitation scenario using the BEM-FMM method for the entire combined boundary element model cosnsided as a whole. The solution obtained predicted a much smaller activating function error. This observation needs to be investigated further.

## Acknowledgements

The authors wish to thank Dr. Aapo Nummenmaa and Dr. Bruce Rosen at the Martinos Center, Massachusetts General Hospital, Boston for insight and support, and Dr. Leslie Greengard of the Courant Institute of Mathematical Sciences, New York, NY for useful comments. This work was supported in part by the National Institutes of Health Grants R01MH111829 and P41EB015896.

